# An easy estimate of the PFDD for a plant illuminated with white LEDs: 1000 lx = 15 μmol/s/m^2^

**DOI:** 10.1101/289280

**Authors:** Anton Sharakshane

## Abstract

Dependencies have been shown and conversion factors have been determined, which allow to estimate PPFD, YPFD and radiometric power density of white LED light according to the known illumination in lux. A technique for estimating photosynthetically active radiation, which has an adequate accuracy for the task of illuminating plants, has been determined.

## Introduction

### Need for white light

Pink light with a depleted spectrum or yellow light of a sodium lamp (HPS) can be used in industrial plant growing. White light is necessary when people share a common room with plants and when the decorative qualities of a plant are important. For example, in education projects, plants should be constantly observable. There is usually no separate room for them. Therefore, in education projects, there is also no alternative to white light of high colour rendering, which provide visual comfort for a person and good conditions for plant development [1].

### Measuring and recording the lighting system parameters

For example one growing system built by children, has a complete set of lighting parameters: {0.3 m^2^; 50 W; 11,000 lx; 3,000 К; R_a_ = 98; 165 μmol/s/m^2^; 24×7}. The parameters may be suboptimal, but their registration allows to discuss, adopt practices, offer and try other options. Not to make such a registry in the education project is incorrect and unpedagogical.

To estimate the intensity of plant illumination with non-white light, a spectrometer is required. The illumination with white light is measured by a luxmeter. And since the white light spectrum profile is usually described by the known colour temperature and colour rendition with sufficient accuracy for agrotechnical purposes [1], measuring the illumination in lux allows to estimate photosynthetically active radiation in any other units used in growing plants.

### Comparative effectiveness of using white light

The effectiveness of modern white LED luminaires, expressed in μmol/J in the relevant range of 400…700 nm, nearly corresponds to the best specialized HPS and is slightly inferior to LED luminaires with a depleted spectrum [1]. This makes the usage of white light energetically reasonable.

Based upon the data in [2], specialized HPS for greenhouse lighting with the power of 600…1,000 W have an effectiveness of about 1.6 μmol/J; 1,000 lm of luminous flux correspond to about PPF = 12 μmol/s, and the illumination of 1,000 lx corresponds to about PPFD = 12 μmol/s/m^2^. Knowing the rules for converting luminous flux units for white LED light, the experience of lighting plants with HPS lamps can be used in industrial greenhouses.

### Comparison of different spectra options for lighting plants

A direct comparison of the spectra of light sources (Fig. 1) shows that the light of white LEDs with the most common parameters 4,000K and R_a_=80 is richer than the HPS spectrum and somewhat inferior (in terms of the red component content) to a typical spectrum of pink light for illuminating plants with an accustomed but obviously incorrect commercial name ‘grow light full spectrum’. White light with a high colour rendering index is richer in the spectral content than other options and closer to the continuous spectrum of natural light.

**Fig. 1.**
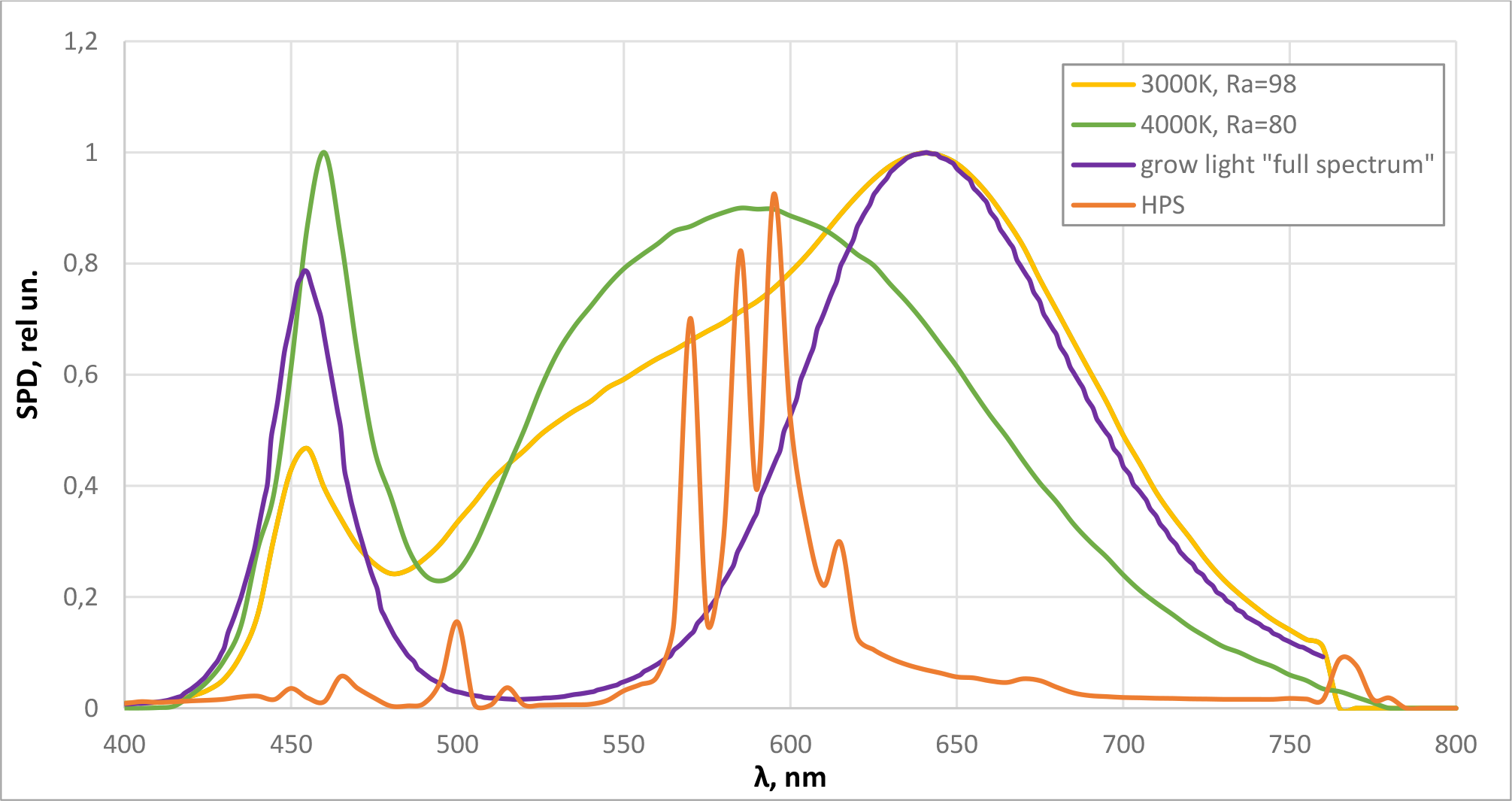
Comparing the spectra of white LED light and specialized light for growing plants

The curves show that the growth of white light colour rendering leads to an increase in the proportion of light useless for photosynthesis with a wavelength greater than 700 nm. But this proportion does not exceed a few percent and not higher than that of the ‘grow light full spectrum’.

Spectral components that perform only a signal function and are not included in the white LED light spectrum (primarily 400 nm and 730 nm) can be added to white light using separate luminaires with narrowband LEDs at this wavelength. The experimental test of the expediency of such adding and the determination of its optimum intensity for each crop is quite simple. But the basic energy demand of a plant for light is to be met first.

### Objective

The work objective is to analyze the accuracy of estimates of various parameters of plant lighting with white light, to compare the errors introduced by various factors, and to determine the methodology with adequate for practical problems accuracy.

## Methods

The calculations were carried out based on the same database of 205 spectra of white LEDs, which was used in the works [1, 2]. We carried out a multivariate regression analysis using models involving independent variables raised to from the negative 1st to the 2nd power and searched for simplified models where the magnitude of error is slightly higher than when using a complete model.

## Results

### LER: Luminaire Efficacy Rating

The parameter LER [lm/W] has the same dimension as the luminous efficacy η [lm/W] characterizing the luminaire, but it denotes the luminous flux in lumens corresponding to one watt of radiometric radiation power.

The LER is slightly dependent on the colour temperature CCT and has a significant spread with a constant colour rendering Ra (Fig. 2). But for estimation, we can use the formula obtained by the regression analysis LER=-196.4+14.5⋅Ra-0.1⋅Ra^2^ at δ=2.7%.

**Fig. 2.**
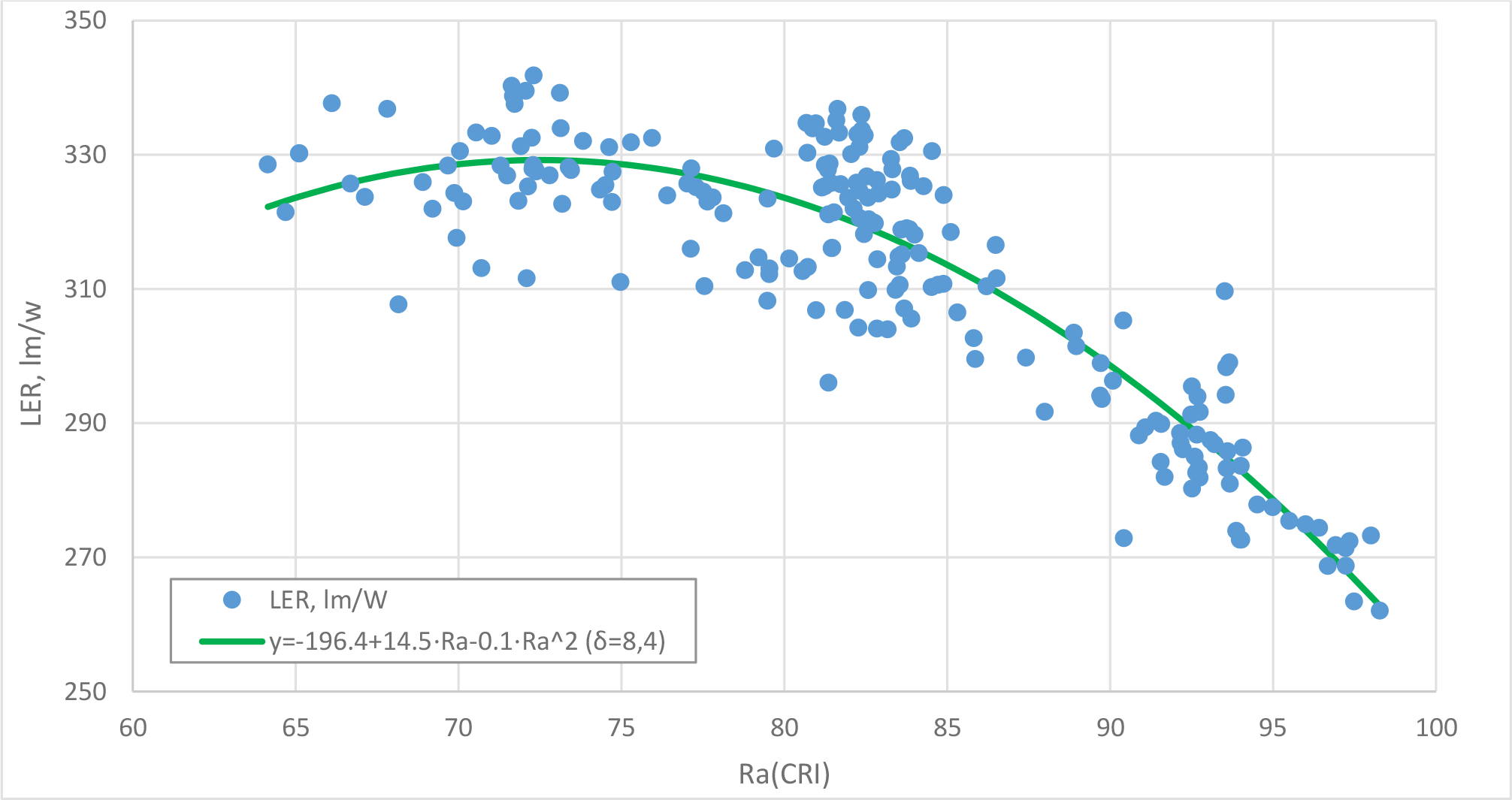
Dependence of the LER of white LED light on the overall colour rendering index

If we use the round value LER = 300 lm/W to estimate the LER, the standard deviation will be 7.3% of the estimated value.

Knowing the LER value, it is easy to calculate the radiometric power according to the formula W = F / LER and the radiometric power density W / S = E / LER, where W[W] – radiometric power, F[lm] – luminous flux, S[m^2^] – area where the luminous flux falls, E[lx] – illumination.

The radiometric density of a luminous flux is rarely used in plant lighting guidelines. The LER estimate is useful in understanding that the radiometric flux density is proportional to the illumination in lux, and the spectral parameters of white light can be neglected in a first approximation. The LER estimate also makes it possible to estimate the efficiency of a lighting unit in the large using the formula η = 100 · E ·S / LER / P, where E[lx] – the actual measured illumination created on the area S[m^2^] by the lighting unit consuming the power P[W]. Efficiency is an important integral parameter of effectiveness monitoring.

### Energy value of a unit of light

The energy value of light for a plant in a first approximation is determined by the value of PPF (Photosynthetic Photon Flux) in micromoles per second in the range of 400-700 nm, or more accurately by the value of YPF (Yield Photon Flux) corrected for the McCree (1972) curve [3].

Using the value of YPF instead of PPF (or in terms of the illuminated area, YPFD instead of PPFD) allows to estimate the energy value of light more accurately and choose a light source. Most of the data in the scientific literature to be relied on when estimating the lighting system handles the values of PPF, and this makes the PPF-YPF ratio analysis interesting.

For white light, the PPF-YPF dependence is rather tight, slightly depends on the colour rendering and is determined by the colour temperature (Fig. 3). The regression analysis gives the following expression: PPF / YPF = 1.0045+4.15⋅10^−5^⋅CCT-2.4⋅10^−9^⋅CCT^2^ at δ=0.3%.

**Fig. 3.**
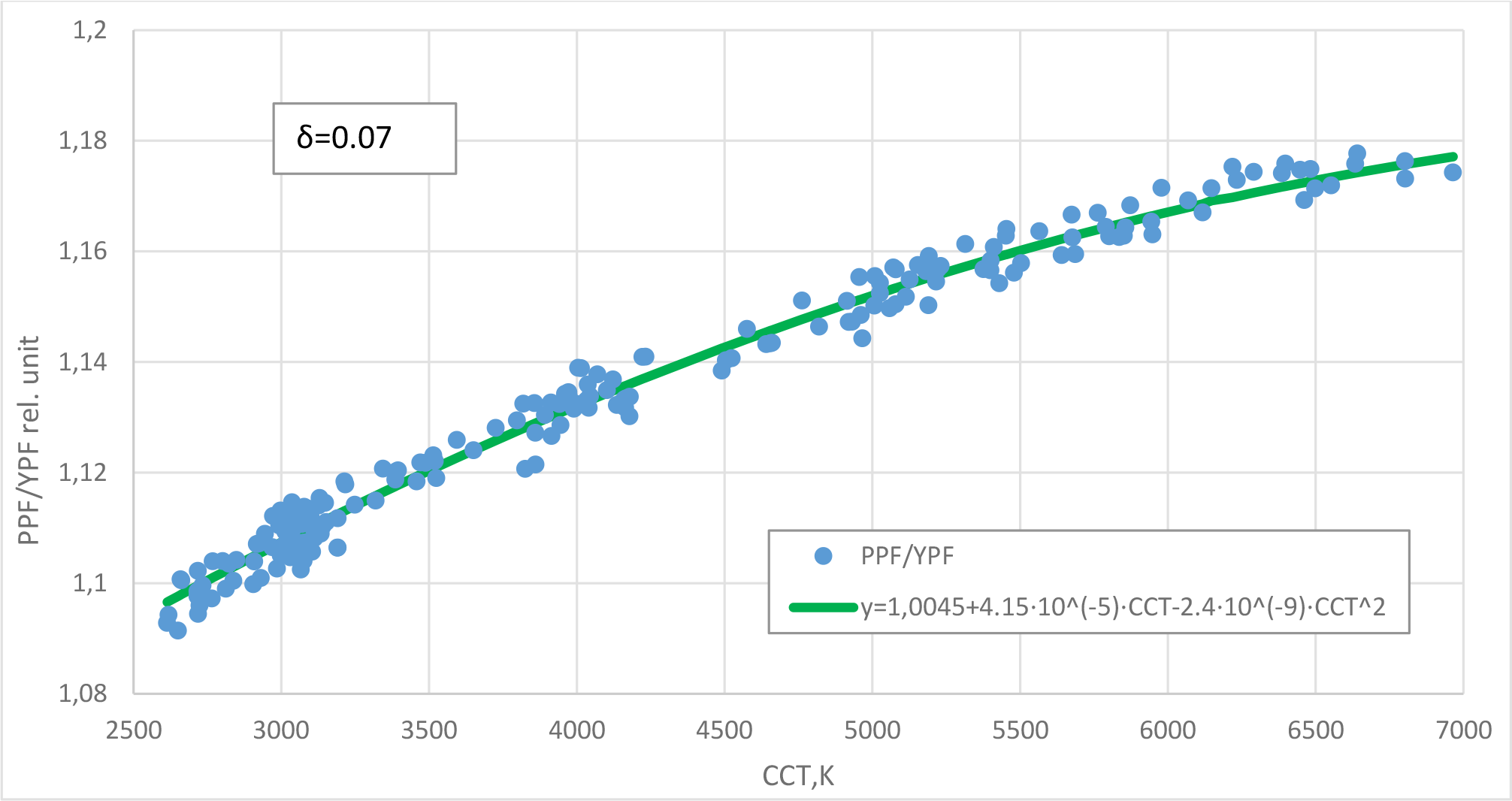
Dependence of the PPF-YPF ratio on the colour temperature of white light

For practical purposes, it is sufficient to take into account that the dependence is almost linear, and for 3,000 K, the PPF is greater than YPF by approximately 10%, and for 5,000 K by 15%. It means about 5% greater energy value of warm light for a plant compared to cold light with equal illumination in lux.

### PPF & PPFD

The regression analysis gives the following PPF estimate corresponding to a luminous flux of 1,000 lm:

PPF{μmol/s/m^2^/klm} = −48+0.0005⋅CCT+12,000/CCT+0.4⋅Ra+2,000/Ra, at δ=2.1%.

For typical values of the spectral parameters, the PPF and PPFD are the following:

| **PPF in $\mu\text{mol/s}$ per 1,000 lm (or PPFD in $\mu\text{mol/s/m}^2$ per 1,000 lx)** |  |  |  |
| --- | --- | --- | --- |
| **CCT \ $R_a$** | **$R_a = 70$** | **$R_a = 80$** | **$R_a = 90$** |
| **3,000 K** | 14.1 | 14.5 | 15.7 |
| **4,000 K** | 13.6 | 14.0 | 15.2 |
| **5,000 K** | 13.5 | 13.9 | 15.1 |

As can be seen from the given values, warm light and light with high colour rendering have a slightly greater energy value for a plant with equal illumination.

The values in the table differ from the round value of 15 units by no more than 7%, so for practical purposes, the following rule can be used: <u>the flow of 1,000 lm corresponds to</u> PPF = 15 μmol/s, and the illumination of 1,000 lx corresponds to PPFD = 15 μmol/s/m^2^.

### Estimating the luminous flux utilization factor

The luminous flux utilization factor *k* is a fraction of the luminous flux from the lighting unit falling on the leaves of plants. This value can be used, for example, to estimate PPFD in accordance with the above dependence by the formula: PPFD[μmol/s/m^2^] = k·15·F[klm]/S[m^2^], where F – luminous flux in kilolumens, S – illuminated area in square meters.

Uncertainty of *k* increases the error in estimating photosynthetically active radiation. Let us consider the possible values of *k* for the main types of lighting systems:

#### 1 Point and line sources

The illumination produced by a point source at the local site decreases in inverse proportion to the squared distance between this site and the source. The illumination produced by extended line sources over narrow beds decreases in inverse proportion to the distance.

The illumination decreases not due to the fact that the light ‘weakens’ with the distance, but due to the fact that as the distance increases, an increasing proportion of light goes past the leaves. This makes it extremely unprofitable to illuminate single plants or single extended beds with highly-suspended luminaires. Optics narrowing the luminous flux allow to send a large proportion of the luminous flux to the plant, but in general, the exact proportion remains unknown.

A strong dependence of illumination on distance and the uncertainty of the effect of using optics do not allow us to determine the utilization factor *k* in the general case.

#### 2 Reflecting surfaces

When using closed volumes with perfectly reflecting walls, the entire luminous flux hits the plant. However, the actual reflection coefficient of mirror or white surfaces is always less than unity. And this leads to the fact that the proportion of the luminous flux reaching the plants still depends on the reflective properties of surfaces and on the volume geometry. And it is impossible to determine *k* in the general case.

#### 3 Large arrays of sources over large planting areas

Large arrays of point or line luminaires over large planting areas are energetically favourable. Quantum radiated in any direction will certainly reach some of the leaves; the factor *k* is close to 1.0.

Intermediate conclusion: for all considered geometries of the lighting unit, the uncertainty of the light fraction reaching the plants is higher than the difference between PPFD and YPFD and higher than the error determined by the unknown colour temperature and colour rendering. Therefore, for a practical evaluation of the intensity of photosynthetically active radiation, it is reasonable to choose a fairly rough illumination estimation technique that does not take into account these peculiarities. And, if possible, measure the actual illumination by a luxmeter.

### Illumination measurement error

In direct measurements, it is necessary to take into account the unevenness of the illumination created by the lighting unit. A case study: the standard EN 12464-1 “The Lighting of Workplaces” requires the ratio of minimum to average illumination of no more than 0.7. What in practice means the difference in illumination of different areas up to 30% and a significant error of the average value with a small number of measurements. In addition, the luxmeter readings may differ from the true values by 10…20% in accordance with the instrument precision as per DIN 5032-7.

### Effect of the error in the PPFD value on the result

In accordance with the constraining factor law (Liebig’s Barrel), the scarce factor (e.g., light) affects the yield linearly. However, the optimum level of PPFD is usually chosen by the yield maximization criterion, and therefore, at the boundary or beyond the boundary of the linear dependence. For example, in [4], the optimum illumination intensity for Chinese cabbage was determined PPFD = 340 μmol/s/m^2^. And the argument that at higher illumination levels the yield is slightly heavier so that an increase in illumination becomes economically impractical was used as the criterion. In a private communication, the authors of this work pointed out that with an improved technique for growing the same crop, a linear increase in yield was observed at illuminances up to 500 μmol/s/m^2^.

Thus, the situation of a significant impact of the PPFD level on crop yield is in itself a sign of insufficient illumination. A sufficient quantity of light negates the significance of the error in determining the illumination level and makes the use of high-precision estimates unreasonable.

## Conclusion

The most adequate estimation of the photosynthetically active white light flux is achieved if one measures the illumination E in kilolumens using a luxmeter, neglects the influence of spectral parameters, and estimates the PPFD of white LED light according to the formula:

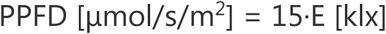

## Acknowledgments

The author gratefully acknowledges the help of Irina O. Konovalova, senior staff scientist of the IBMP RAS, PhD in Biology; Nikolay N. Sleptsov, Gorshkoff.ru chief technology officer; Mikhail Chervinsky, FAE at Cree; Anna G. Savitskaya, lighting technician; Alexander A. Sharakshane, senior staff scientist of the IRE RAS, PhD in Physics and Mathematics; Andrei A. Anosov, leading staff scientist of the IRE RAS and professor at Sechenov MSMU, Doctor of Sciences in Mathematics and Physics.

